## SupplementaryMaterial for "A large-scale resource for tissue-specific CRISPR mutagenesis in *Drosophila*"

**for**

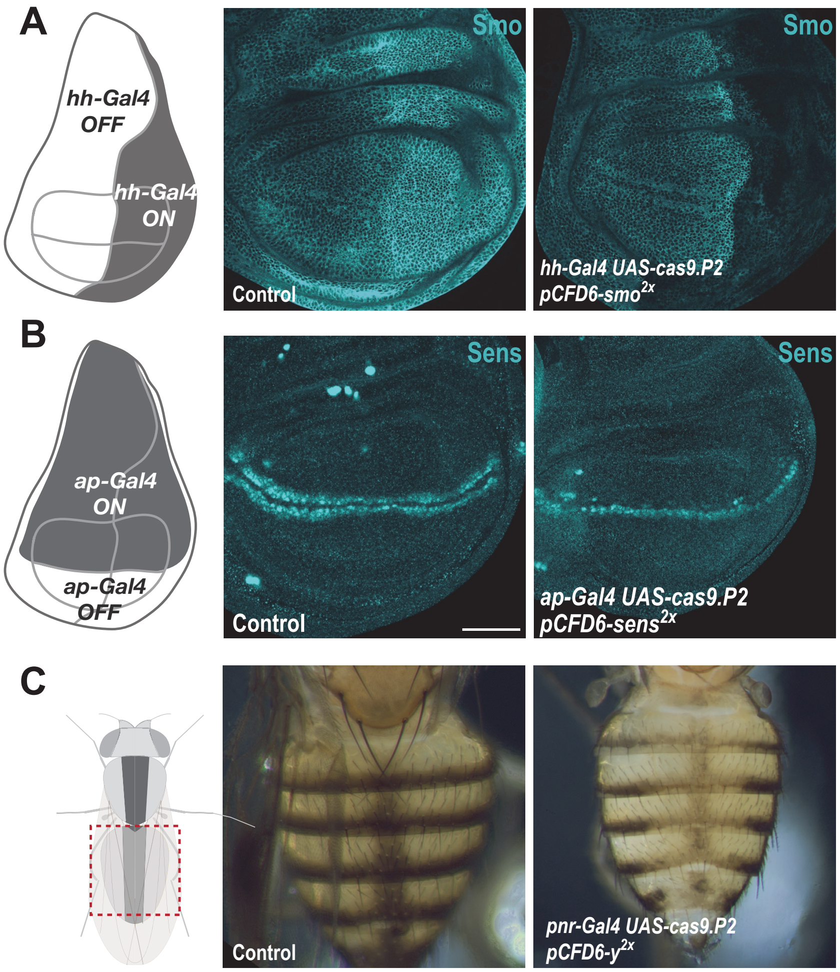


**Supplementary Figure 1: Efficient conditional CRISPR mutagenesis in various *Drosophila* tissues.** (**A**) CRISPR mutagenesis of *smo* in the posterior compartment of the wing imaginal disc. Smo protein was detected by immunohistochemistry. Smo is normally expressed in all wing disc cells, but protein levels are higher in the posterior compartment (see Control (*hh-Gal4 UAS-cas9.P2*)). In *hh-Gal4 UAS-cas9.P2 pCFD6-smo*$^{2x}$ wing disc cells in the posterior compartment express no or reduced levels of Smo, presumably reflecting cells containing only one or no functional *smo* alleles. (**B**) CRISPR mutagenesis of *sens* in the dorsal compartment of wing imaginal discs with *ap-Gal4* leads to a loss of Sens expression in most, but not all cells. (**C**) Mutagenesis of *y* in the dorsal abdomen. In *pnr-Gal4 UAS-cas9.P2 pCFD6-y*$^{2x}$ animals cuticle coloration is uniformly changed in a broad stripe centred around the dorsal midline, compared to control animals (*pnr-Gal4 UAS-cas9.P2 pCFD6-Sfp24C1*$^{2x}$). Note that the strong phenotype mediated by *pCFD6-y*$^{2x}$ is in line with the high levels of mutagenesis with this construct reported in Supplementary Fig. 6.

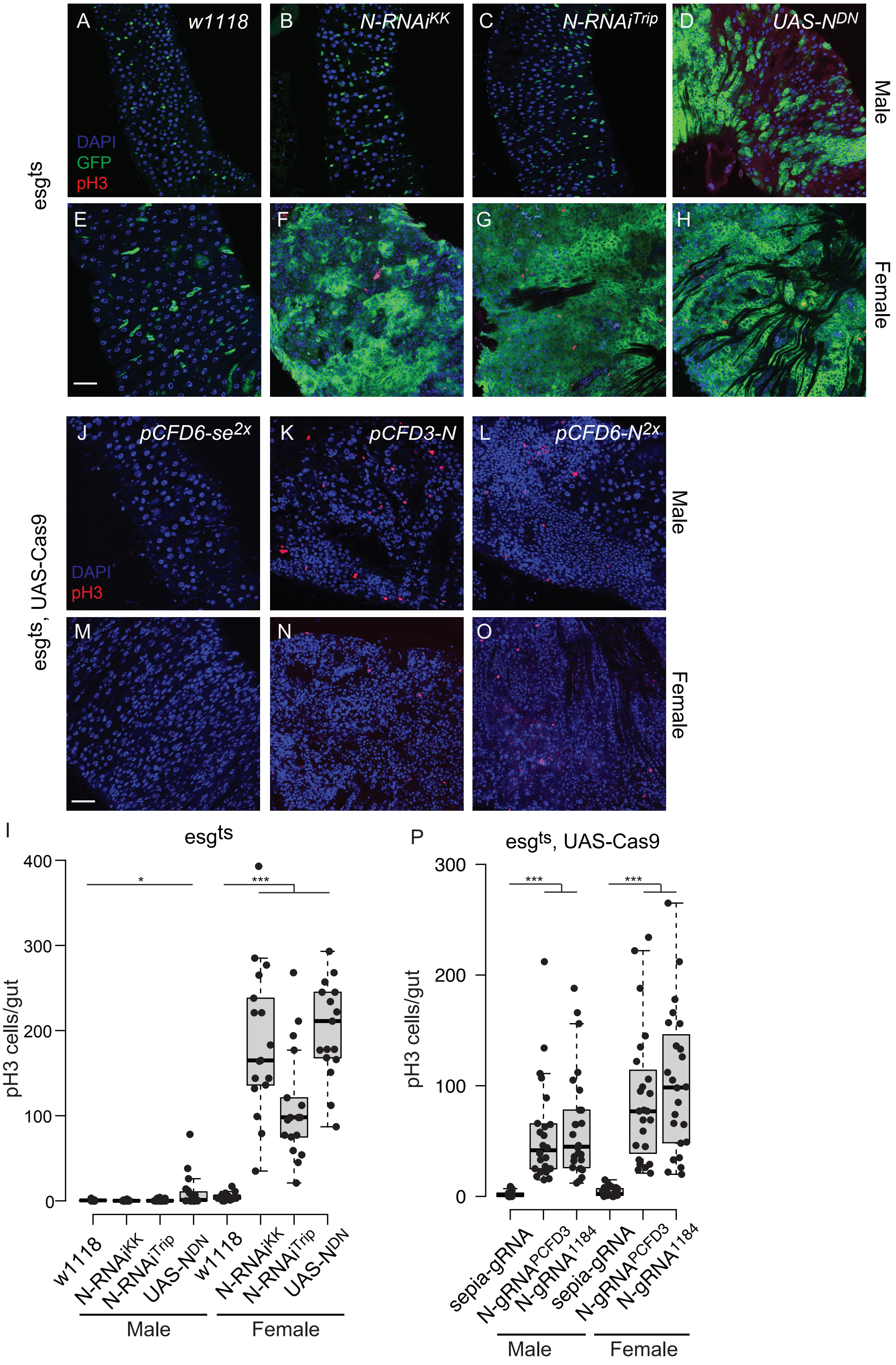

A
w1118
B
N-RNAi^KK
C
N-RNAi^Trip
D
UAS-N^DN
Male
DAPI
GFP
pH3
esgts
E
F
G
H
Female
J
pCFD6-se^2x
K
pCFD3-N
L
pCFD6-N^2x
Male
DAPI
pH3
esgts, UAS-Cas9
M
N
O
Female
I
esgts
pH3 cells/gut
400
300
200
100
0
*
***
w1118
N-RNAi^KK
N-RNAi^Trip
UAS-N^DN
w1118
N-RNAi^KK
N-RNAi^Trip
UAS-N^DN
Male
Female
P
esgts, UAS-Cas9
pH3 cells/gut
300
200
100
0
***
***
sepia-gRNA
N-gRNA^PCFD3
N-gRNA^1184
sepia-gRNA
N-gRNA^PCFD3
N-gRNA^1184
Male
Female

**Supplementary Figure 2: Qualitative differences between CRISPR mutagenesis and RNAi knock-down of *Notch* in the *Drosophila* midgut.** (**A-H**) Representative images of the posterior midgut of male or female *esg*$^{ts}$ *>w1118* (**A, E**), *esg*$^{ts}$*>N-RNAi*$^{KK}$ (**B, F**), *esg*$^{ts}$*>N-RNAi*$^{Trip}$ (**C, G**) and *esg*$^{ts}$*>UAS-N*$^{DN}$ animals, stained with pH3 antibody in red to mark mitotic cells. *esg-GFP* is shown in green, nuclei are stained with DAPI (blue). Animals were raised at 18°C to adulthood and then incubated at 29°C for 15 days. (**I**) Quantification of pH3-positive cells per adult midgut of the indicated genotypes after 15 days at 29°C. (**J-O**) Representative images of the posterior midgut of male or female *esg*$^{ts}$ *UAS-Cas9.P2 pCFD6-se*$^{2x}$ (**J, M**), *esg*$^{ts}$ *UAS-Cas9.P2 pCFD3-N* (**K, N**) and *esg*$^{ts}$ *UAS-Cas9.P2 pCFD6-N*$^{2x}$ (**L, O**) flies stained with pH3 antibody in red and nuclei are stained with DAPI (blue). Animals were raised at 18°C, after eclosion mutagenesis was induced for 5 days at 29°C, followed by 18°C for 15 days. Unlike in the RNAi condition, no GFP is visible due to repression by Gal80 at 18°C. (**P**) Quantification of pH3-positive cells per adult midgut of the indicated genotypes at 18°C for 30 days after inducing CRISPR/Cas9 mediated mutagenesis. Significant differences in the number of pH3-positive cells between Notch RNAi or knock-out group (*esg*$^{ts}$ *> N-RNAi*$^{KK}$*, esg*$^{ts}$ *> N-RNAi*$^{trip}$*, esg*$^{ts}$*;UAS-Cas9 > N-gRNA* $^{PCFD3}$ *or esg*$^{ts}$*;UAS-Cas9 > N-gRNA*$^{1184}$) and the control groups. (*esg*$^{ts}$ *> w1118* or *esg*$^{ts}$*;UAS-Cas9 > sepia-gRNA* ) are indicated with asterisks (* p<0.05; ** p<0.01; *** p<0.001; two-tailed T-test) Scale bars: 30 µm (**A-H**, **J-O**). Note that the total number of mitotic cells in the RNAi and CRISPR conditions cannot be directly compared, as animals were raised at different temperatures.

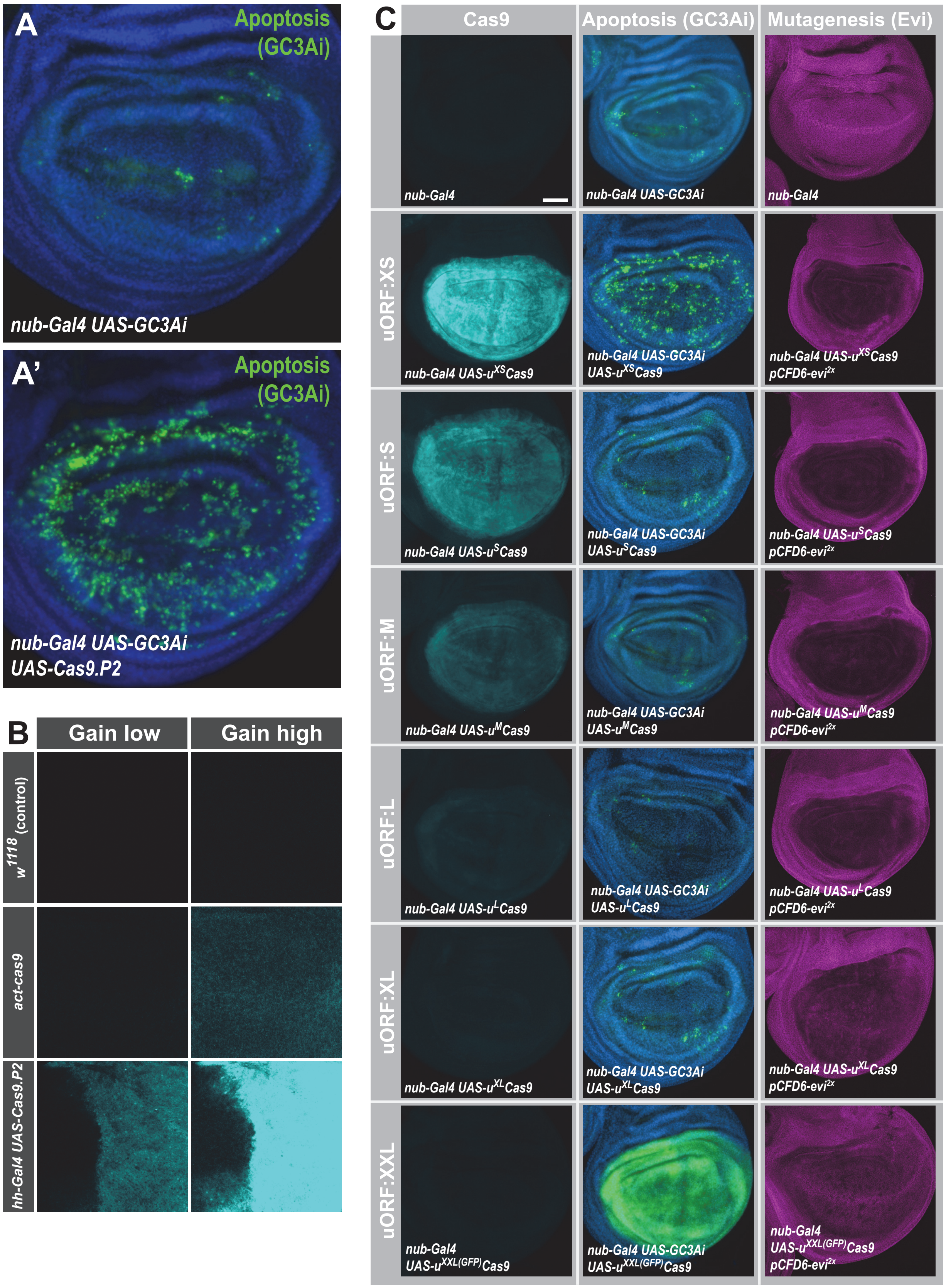
A
Apoptosis (GC3Ai)
nub-Gal4 UAS-GC3Ai
A'
Apoptosis (GC3Ai)
nub-Gal4 UAS-GC3Ai UAS-Cas9.P2
B
Gain low
Gain high
w1118 (control)
act-cas9
hh-Gal4 UAS-Cas9.P2
C
Cas9
Apoptosis (GC3Ai)
Mutagenesis (Evi)
nub-Gal4
nub-Gal4 UAS-GC3Ai
nub-Gal4
uORF:XS
nub-Gal4 UAS-uXSCas9
nub-Gal4 UAS-GC3Ai UAS-uXSCas9
nub-Gal4 UAS-uXSCas9 pCFD6-evi2x
uORF:S
nub-Gal4 UAS-uSCas9
nub-Gal4 UAS-GC3Ai UAS-uSCas9
nub-Gal4 UAS-uSCas9 pCFD6-evi2x
uORF:M
nub-Gal4 UAS-uMCas9
nub-Gal4 UAS-GC3Ai UAS-uMCas9
nub-Gal4 UAS-uMCas9 pCFD6-evi2x
uORF:L
nub-Gal4 UAS-uLCas9
nub-Gal4 UAS-GC3Ai UAS-uLCas9
nub-Gal4 UAS-uLCas9 pCFD6-evi2x
uORF:XL
nub-Gal4 UAS-uXLCas9
nub-Gal4 UAS-GC3Ai UAS-uXLCas9
nub-Gal4 UAS-uXLCas9 pCFD6-evi2x
uORF:XXL
nub-Gal4 UAS-uXXL(GFP)Cas9
nub-Gal4 UAS-GC3Ai UAS-uXXL(GFP)Cas9
nub-Gal4 UAS-uXXL(GFP)Cas9 pCFD6-evi2x

**Supplementary Figure 3: High levels of Cas9 expression from *UAS-cas9.P2* are cytotoxic.** (**A**) Expression of Cas9 in *nub-Gal4 UAS-Cas9.P2* causes excessive cell death. Apoptotic cells were visualized by co-expression of GC3Ai, which is fluorescent in cells with activated caspase 3. Expression of GC3Ai in the wing pouch highlights a few cells undergoing apoptosis. Additional expression of *UAS-cas9.P2* causes a dramatic increase in the cells undergoing programmed cell death, highlighting cytotoxicity caused by high levels of Cas9. (**B**) Comparison of Cas9 expression levels between *act-cas9* and *hh-Gal4 UAS-cas9.P2* wing discs. Cas9 was detected by antibody staining and imaged in the same session with identical setting at either low (top panel) or high (bottom panel) detector gain. The difference in Cas9 levels are such, that Cas9 expressed from *act-cas9* is undetectable with low detector gain and Cas9 staining in *hh-Gal4 UAS-cas9.P2* wing discs is oversaturated with high detector gain, where Cas9 from *act-cas9* is visible. The act-cas9 transgene is known to mediate highly efficient mutagenesis in *Drosophila* (Port et al., 2014; Port et al. 2015). (**C**) Systematic characterization of Cas9 expression, toxicity and mutagenesis of the UAS-uCas9 series. Transgenes of the UAS-uCas9 series were recombined with *nub-Gal4* and wing imaginal discs were stained for Cas9 protein. Cas9 levels gradually reduce as the size of the uORF increases (left panel). *nub-Gal4 UAS-uCas9* flies were crossed to *UAS-GC3Ai* to visualize cells undergoing apoptosis. Elevated levels of apoptosis were only observed with *UAS-*$u^{XS}$*Cas9*. The longest uORF ($u^{XXL}$) encodes EGFP, preventing visualization of dying cells with GC3Ai (middle panel). *nub-Gal4 UAS-uCas9* flies were crossed to *pCFD6-evi*$^{2x}$ and mutagenesis of *evi* was indirectly observed by loss of Evi staining. All transgenes of the UAS-uCas9 series mediate *evi* mutagenesis, with transgenes containing the four shortest uORFs (XS-L) leading to comparable gene editing that removes Evi from nearly all cells in the Gal4 expression domain (right panel).

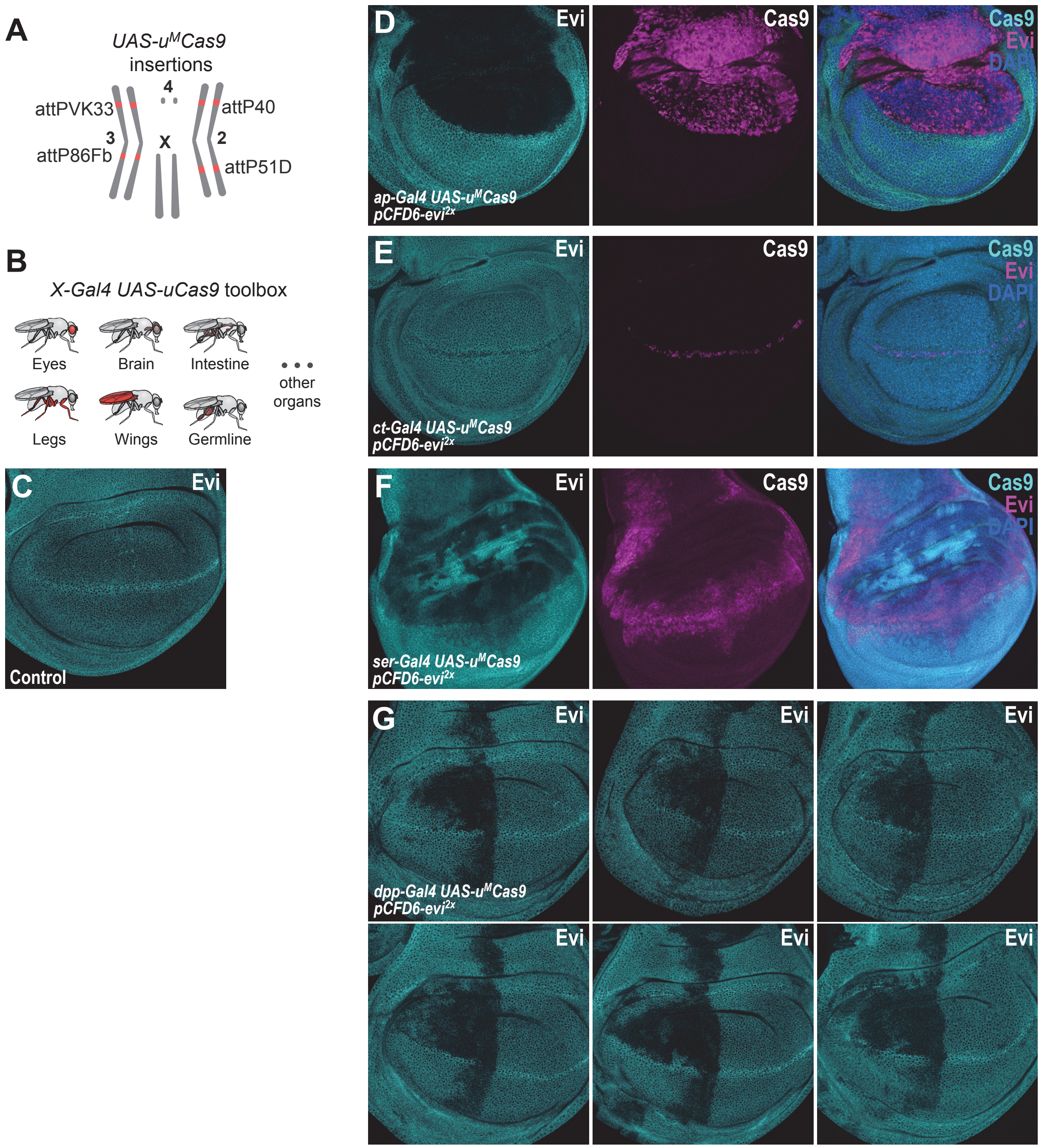


**Supplementary Figure 4: CRISPR mutagenesis patterns reflect expression patterns of Gal4 lines throughout develpment.** (**A**) Chromosomal locations of *UAS-u$^{M}$Cas9* transgenes currently available. (**B**) A growing collection of *Gal4 UAS-uCas9* stocks will allow tissue specific mutagenesis in *Drosophila*. (**C**) Evi is expressed in all cells of wildtype third-instar wing imaginal discs and accumulates in cells at the dorsal-ventral boundary. (**D**) Mutagenesis of *evi* in *ap-Gal4 UAS-u$^{M}$Cas9 pCFD6-evi$^{2x}$* wing discs results in loss of Evi protein exclusively in the dorsal compartment, which expresses Cas9 protein. (**E**) CRISPR mutagenesis of Evi with *ct-Gal4* results in loss of Evi staining only in cells along the dorsal-ventral boundary, where also Cas9 protein is expressed. (**F**) CRISPR mutagenesis with ser-Gal4 results in loss of Evi staining in the pouch, hinge and notum region of the wing disc and includes large areas where no Cas9 protein is detectable in third instar wing discs. (**G**) Mutagenesis of *evi* with *dpp-Gal4*. Dpp is known to be expressed in a stripe along the anterior-posterior boundary in third instar wing imaginal discs. In addition to this domain, mutagenesis is consistently found in the anterior-dorsal region of the pouch. Six representative discs for the same genotype are shown.

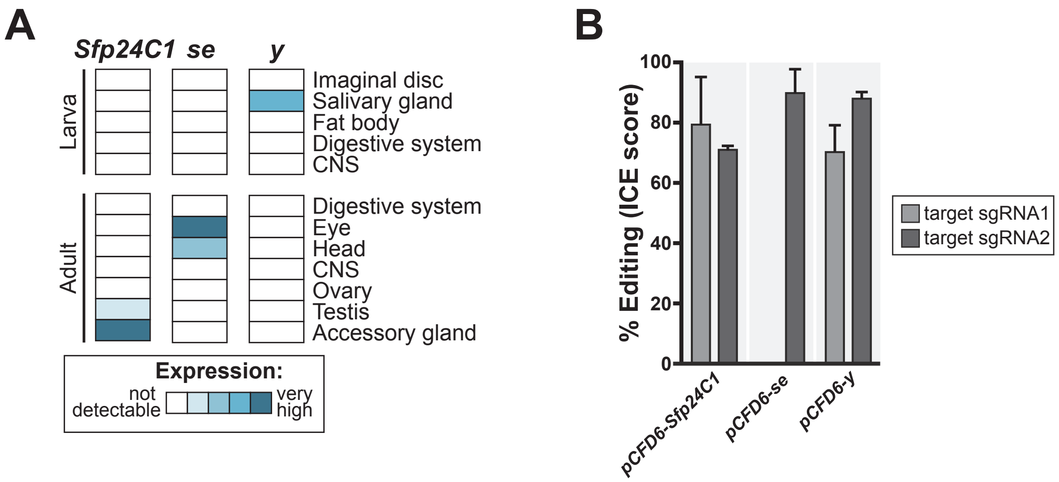


**Supplementary Fig. 5: Negative control sgRNA transgenes for use with HD_CFD library.** (**A**) *Sfp24C1*, *se* and *y* are not expressed in most tissues (upper panel, data from modENCODE), making them suitable as negative controls in most circumstances. (**B**) sgRNA lines mediate efficient mutagenesis at both (*pCFD6-Sfp24C1*$^{2x}$, *pCFD6-y*$^{2x}$) or one (*pCFD6-se*$^{2x}$) target sites.

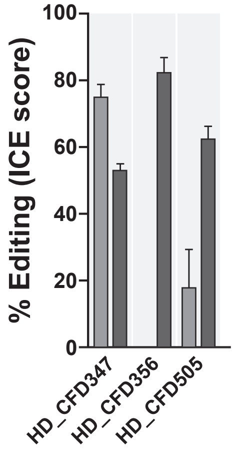


**Supplementary Figure 6: HD_CFD sgRNA line resulting in false-negative results mediate efficient on-target mutagenesis.** Lines HD_CFD347 (Target: *hbn*), HD_CFD356 (Targat: *Ets97D*) and HD_CFD505 (Target: *gcm*) give rise to viable offspring when crossed to *act-*

*cas9;;tub-Gal4/TM6B*. Failure to result in lethality does not reflect inactive sgRNAs. Target locus was amplified by PCR and editing efficiency was measured by Sanger sequencing followed by ICE analysis. Bars represent mean and error bars show standard deviation.
